# Phylogenetic analysis of SARS-CoV-2 data is difficult

**DOI:** 10.1101/2020.08.05.239046

**Authors:** Benoit Morel, Pierre Barbera, Lucas Czech, Ben Bettisworth, Lukas Hübner, Sarah Lutteropp, Dora Serdari, Evangelia-Georgia Kostaki, Ioannis Mamais, Alexey M Kozlov, Pavlos Pavlidis, Dimitrios Paraskevis, Alexandros Stamatakis

## Abstract

Numerous studies covering some aspects of SARS-CoV-2 data analyses are being published on a daily basis, including a regularly updated phylogeny on nextstrain.org. Here, we review the difficulties of inferring reliable phylogenies by example of a data snapshot comprising all virus sequences available on May 5, 2020 from gisaid.org. We find that it is difficult to infer a reliable phylogeny on these data due to the large number of sequences in conjunction with the low number of mutations. We further find that rooting the inferred phylogeny with some degree of confidence either via the bat and pangolin outgroups or by applying novel computational methods on the ingroup phylogeny does not appear to be possible. Finally, an automatic classification of the current sequences into sub-classes based on statistical criteria is also not possible, as the sequences are too closely related. We conclude that, although the application of phylogenetic methods to disentangle the evolution and spread of COVID-19 provides some insight, results of phylogenetic analyses, in particular those conducted under the default settings of current phylogenetic inference tools, as well as downstream analyses on the inferred phylogenies, should be considered and interpreted with extreme caution.

## Introduction

The Coronavirus disease 2019 (COVID-2019) caused by a novel coronavirus [severe acute respiratory syndrome coronavirus-2 (SARS-CoV-2)] emerged in Wuhan, China in December 2019 (WHO situation report May 30, 2020) and spread worldwide in 212 countries and territories causing more than 17.6 million cases and 680,000 deaths within a period of 7 months (WHO situation report August 2, 2020).

A full genome sequence analysis revealed that 2019-nCoV-2 belongs to the betacoronaviruses, but that it is divergent from the SARS-CoV and MERS-CoV that caused past epidemics. The 2019-nCoV-2 and the bat-SARS-like coronavirus form a distinct lineage within the subgenus of the Sarbecovirus. The whole-genome sequence of SaRS-CoV-2 shows 96.2% similarity to that of a bat SARS-related coronavirus (RaTG13) collected in the Yunnan province of China. The SARS-CoV-2 is also closely related to the coronavirus from Malayan pangolin in a particular genomic region coding for the spike protein, including the receptor-binding domain. This is notable, because the remainder of the viral genome is most closely related to the bat coronavirus RaTG13 (1). The latter observation suggests a putative recombination event between viruses infecting bats, pangolins, and humans.

Since the early characterization of SARS-CoV-2 in Hubei, China, an enormous number of sequences have been characterized. On July 31st 2020, approximately 75,000 full genome sequences have become available. Molecular epidemiology has attempted to provide a detailed picture about the distinct lineages and sub-strains circulating in different geographic areas as well as about the dispersal pattern and cross-border transmissions at different time periods during the pandemic (https://nextstrain.org/ncov/global; (2)). Moreover, whole-genome sequence analysis has been used for within-country studies as well as for the detailed investigation of viral dispersal within specific communities.

To date, the globally circulating viruses have been classified into 6 major clades denoted as *S, L, V, G, GH*, and *GR* (https://www.gisaid.org/ (3)). Analyses of the viral sequences can unravel the number of mutations separating the lineages from the founding Wuhan haplotype. These analyses provide a more detailed classification of variants into haplotypes that can be used to trace the geographical distribution and patterns of dispersal of distinct lineages (4). Molecular epidemiology studies attempt to quantify the number of introductions of SARS-CoV-2 to different countries and their putative geographic origin (5, 6). Haplotype analyses based on SARS-CoV-2 can also provide information about within-country infection clusters.

An analysis from Iceland applied molecular analyses and verified their results via a comparison with contact tracing networks (7). It concludes that, when contact tracing networks are unavailable, phylogenetic analyses can be deployed to disentangle infection clusters within countries. Full genome analyses of SARS-CoV-2 can potentially identify emerging novel variants that may alter the spike interaction with the ACE2 receptor, TMPRSS2 protease, and epitope mapping. This has been previously shown in a study by Korber *et al*. (8), suggesting that viruses harboring the D614G mutation were associated with increased SARS-CoV-2 viral loads, yet not associated with increased disease severity. These findings were also supported by Bedford’s group (the Bedford lab in Washington: https://github.com/blab/ncov-D614G).

SARS-CoV-2 evolves at an estimated nucleotide substitution rate ranging between 10^−3^ and 10^−4^ substitutions per site and per year (see Table 1 in (6)). Molecular clock analyses have been used to estimate the time of the most recent common ancestor (MRCA) of the global pandemic as well as the MRCA of local epidemics in different geographical regions (see again Table 1 in (6)).

**Table 1.** Log likelihood scores of the best-scoring ML tree topology (FMSA-B) after model parameter (`GTR`, ML base frequencies and Γ rate heterogeneity) and branch length optimization with the following (default) settings: `blmax`: 100, fast branch length optimization, `lh`_*ϵ*_: 0.1, and varying the indicated `brmin` (* default) value.

| **blmin** | **LLH RAXML-NG** | **LLH IQ-Tree** |
| --- | --- | --- |
| 1e−1 | −65048.221 | −65048.222 |
| 1e−2 | −65048.221 | −65048.222 |
| 1e−3 | −65037.286 | −65048.222 |
| 1e−4 | −64641.947 | −65048.222 |
| 1e−5 | −64271.858 | −64748.599 |
| 1e−6 | *−64271.716 | *−64310.564 |
| 1e−7 | −64271.706 | −64275.513 |
| 1e−8 | −64271.706 | −64272.022 |
| 1e−9 | −64271.705 | −64271.673 |
| 1e−10 | −64271.671 | −64271.642 |

The inference of a phylogenetic tree on the full genomes is pivotal to numerous molecular epidemiology tools and studies (e.g., (9–13)). A plethora of studies (e.g., (14–22)) to disentangle the evolution of the SARS-CoV-2 pandemic is currently being published at a high pace and under considerable time pressure, both with respect to the tree inference time, paper writing time as well as the review time for these papers. In almost all cases, including the daily updated virus phylogenies on the exceptional nextstrain platform, phylogenetic inference on the currently available virus genomes is conducted predominantly via standard Maximum Likelihood (ML) based tools using default program parameters. In addition, several publications also deploy some of the fast, yet less accurate bootstrapping and tree search options implemented in tools such as standard RAxML (23) and IQ-Tree (24). In general, some of these analyses might have been (too?) rushed, not only at the phylogenetic inference level, but also potentially at previous stages of SARS-CoV-2 related data generation and data analysis steps (e.g., see http://virological.org/t/issues-with-sars-cov-2-sequencing-data/473).

In our study, we do not follow this trend, but take a detailed look at the general difficulties of inferring and post-analyzing phylogenies on the highly challenging SARS-CoV-2 dataset, as it contains thousands of taxa with few mutations, and hence comparatively weak signal. Together, these afore-mentioned difficulties render phylogenetic analysis and post-analysis highly challenging, both with respect to the signal that we can extract, but also regarding the numerical stability of current tools.

The remainder of this paper is structured as follows. We first provide an overview of our data preparation and analysis pipeline. Subsequently, we discuss some noteworthy difficulties that arose when processing the data. Then, we present the results of our inference, rooting, and classification attempts. We conclude the paper with a critical discussion of the results.

## Data Preparation and Analysis Pipeline

Our data analysis pipeline is available at https://github.com/BenoitMorel/covid19_cme_analysis under GNU GPL.

### Raw data pre-processing

We downloaded the raw data from gisaid.org on May 5, 2020. It contained 16,453 full genome (> 29,000 bp) raw sequences with high coverage. High-coverage sequences are defined by GISAID as sequences containing less than 1% Ns (undetermined characters), less than 0.05% unique amino acid mutations, and no insertions/deletions unless these have been verified by the submitter.

### Filtering

We applied additional filters to identify sequences of high quality. We initially removed approximately 7,717 sequences using the following two-step strategy. First, we trimmed external undetermined characters at the beginning and at the end of the genomes (step 1). After this trimming, we only kept sequences with less than 10 internal undetermined characters (step 2). We trimmed external undetermined characters (step 1) prior to filtering (step 2) because our goal was to only use sequences with a low number of internal undetermined characters. External undetermined characters do not affect the alignment quality since not all sequences start at exactly the same nucleotide. The final filtered raw sequence dataset comprised 8,736 SARS-CoV-2 genomes and two outgroup sequences: the bat CoV (hCoV-19/bat/Yunnan/RaTG13/2013; Accession ID EPI_ISL_402131) and pangolin CoV (hCoV-19/pangolin/Guangdong/1/2019; Accession ID: EPI_ISL_410721) genomes.

### Multiple Sequence Alignment

We aligned the 8,736 and 8,738 (including outgroups) trimmed sequences using the parallel version of MAFFT (v.7.205 (25)) with 40 threads.

### Trimming after alignment

After the MSA process, we further trimmed the first and the last 1,000 alignment sites. We applied this additional trimming as the sequencing did not start/finish at the same position for all sequences. Thus, the initial untrimmed MSA was characterized by a large amount of missing data at the beginning and the end of the MSA (see supplementary Figure 6).

Overall, we generated 4 distinct versions of the alignment:

1. A comprehensive (comprising all 8,738 sequences) Full MSA with bat and pangolin Outgroups (FMSAO)
2. A comprehensive Full MSA of 8,736 sequences *without* outgroups (FMSA)
3. A non-comprehensive (not comprising all sites, and, as a consequence of additional removed sequence duplicates, not containing all 8,736 virus sequences; see below) singletons-removed MSA with bat and pangolin Outgroups (SMSAO)
4. A non-comprehensive singletons-removed MSA without outgroups (SMSA)

We generated the non-comprehensive SMSAs by removing so-called singleton sites from the corresponding full MSAs (FMSA, FMSAO). For biallelic sites, that is, sites with only two states, a singleton site is a column of the MSA where the allele with the lowest frequency is only present in but a single sequence (e.g., AAAAAAAAT). Such sites only have a negligible contribution to the tree inference process due to weak phylogenetic signal. Furthermore, singleton sites can represent sequencing errors, as it is expected that most sequencing errors will appear as singleton sites.

The FMSA consists of 3752 polymorphic sites, out of which 2503 are either biallelic singletons (e.g., AAAAAAACAAA) or ‘multi-allelic singletons’ (e.g., AAAAAAACAAG), that is, sites where more than one allele only occurs once, while the site itself is *not* biallelic. Further, we also removed 97 ‘pseudo-singleton’ sites (e.g., AAAAACCCAAG)), that is, sites that are neither biallelic nor multi-allelic singletons, but do contain a nucleotide with only a single occurrence. In our example, G appears only once. The numbers of polymorphic sites, biallelic, and multi-allelic singletons are exactly identical for the FMSAO dataset (we double checked). Even though multi-allelic singletons as well as pseudo-singletons do contain some phylogenetic signal, we decided to remove them as they may also represent sequencing errors. Further, the pseudo-singleton sites only account for a small proportion (about 2.5%) of the overall polymorphic sites.

Our singleton removal strategy is further justified by a recent study that has shown that lab-specific sequencing practices yield mutations that have been observed predominantly or exclusively by single labs. These can in turn affect the phylogeny reconstruction process (26). The authors provide regularly updated masking recommendations at https://github.com/W-L/ProblematicSites_SARS-CoV2/blob/master/problematic_sites_sarsCov2.vcf.

Finally, we removed all duplicate sequences from all of the above input MSAs, that is, all sequences that are exactly identical. We did this because identical sequences do not yield any additional signal for a phylogenetic analysis. Furthermore, duplicate sequences confound the calculation of support values, branch lengths, and needlessly increase the computational cost of the analyses.

After removal of duplicate sequences, the MSAs contained the following number of taxa: FMSAO (4,871), FMSA (4,869), SMSAO (2,904), SMSA (2,888). Note that, the difference in the number of taxa between SMSAO and SMSA is not simply two (i.e., the two outgroups) as conducting the alignment step with and without outgroups yields distinct MSAs that in turn induce a distinct number of identical sequences *after* trimming and singleton removal.

Also note that, when only considering the ingroup alignments, the datasets comprise a low proportion of unique alignment site patterns relative to the genome length: FMSAO/FMSA 4,997, SMSAO 1,781, SMSA 1,679. The number of unique patterns in FMSAO/FMSA is higher than the number of polymorphic sites that we reported previously, as the tool we used to analyze polymorphic sites ignores sites containing gaps. These low overall numbers in conjunction with the high number of taxa already indicate that the phylogenetic analysis is challenging.

### Phylogenetic Inference

We initially determined the best fit model for the data on an earlier sequence snapshot from April 29th using ModelTest-NG (27). ModelTest-NG selected GTR+R4 (GTR model with 4 discrete free rate categories, also frequently referred to as ’free rates model’) as best fit model. The free rates model (R4) exhibits some substantial intrinsic numerical difficulties (see ‘Difficulties’ Section for a more thorough discussion). To this end, we decided to use the numerically more stable GTR+Γ model with 4 discrete rates for all subsequent inferences.

With respect to the tree search strategy *per se*, we first executed 100 RAxML-NG (28) tree searches on an earlier snapshot of the full MSA using 50 randomized stepwise addition order parsimony starting trees and 50 random starting trees. We did this to explore the behavior of tree searches on these data. We observed that tree searches initiated on parsimony starting trees yielded phylogenies with better log likelihood scores (consistently > 1000 log likelihood units). Thus, we executed all subsequent phylogenetic tree searches on the data snapshot of May 5 using parsimony starting trees only.

Moreover, initial analyses of earlier snapshots of the data unsurprisingly showed low bootstrap support values, low transfer bootstrap support values (29), and a phenomenon that we have termed ‘rugged likelihood surface’ (30).

We had already observed such a rugged likelihood surfaces for difficult-to-analyze datasets with few sites and many taxa that do typically not contain strong phylogenetic signal on bacterial 16S datasets before (30). Characteristic of such datasets is that, for instance, 100 independent tree searches for the best-scoring Maximum Likelihood (ML) tree on the original alignment will yield 100 distinct tree topologies with similar likelihood scores. Moreover, as we will show here, most of these ML tree topologies also exhibit a large pairwise topological distance. However, at the same time, we can not deploy standard statistical significance tests to distinguish and select among those topologically diverse trees. This is because most of the resulting trees will not be statistically significantly different from each other with respect to their likelihood scores. Hence, given these substantial uncertainties in the search for the best-scoring ML tree in conjunction with the low number of variable sites we apply the following procedure in an attempt to infer a representative phylogeny:

1. Conduct 100 ML tree searches using ParGenes (31) that seamlessly orchestrates such searches using RAxML-NG from randomized stepwise addition order parsimony trees
2. Apply all statistical significance tests implemented in IQ-Tree to this set of 100 ML trees
3. Assign ML trees to a ‘plausible’ ML tree set that are not significantly worse than the best-scoring ML tree under *any* statistical significance test implemented in IQ-Tree (i.e., this assignment is conservative).
4. Build a majority rule (MR) and an extended majority rule (MRE) consensus tree from the plausible ML tree set. Note that, neither the majority rule, nor the extended majority rule consensus trees will necessarily be strictly bifurcating.

We conducted tree inferences exclusively on the ingroup MSAs (FMSA/SMSA and FMSAO/SMSAO with outgroups removed *after* the alignment process) as the usage of outgroups, in particular, if they are distant from the ingroup as is the case here, can perturb a phylogenetic analysis (32, 33). We describe later on how we place the outgroups onto the ingroup phylogenies *after* the and independently of the ingroup tree inference.

While we believe that building a consensus tree from the plausible ML tree set constitutes a reasonable approach, the fact of having a (in most cases) multifurcating (e.g., MR- or MRE-based) reference tree topology complicates matters for some of the downstream phylogenetic post-analysis methods, which often expect a strictly bifurcating phylogeny as input. The general strategy we adopt for addressing this issue is that, whenever possible, we attempt to compute summary statistics of post-analyses over the individual bifurcating trees in the plausible tree set. This approach is, in a sense, analogous to summarizing a posterior tree set as obtained from Bayesian analyses.

For improved clarity and readability we introduce the following notation for the inferred trees:

- FMSA-C: Majority rule consensus of plausible ML ingroup tree set on FMSA
- FMSA-CE: Extended majority rule consensus of plausible ML ingroup tree set on FMSA
- FMSA-P: Plausible ML tree set for FMSA
- FMSA-B: Best-scoring ML ingroup tree inferred on FMSA
- SMSA-C: Majority rule consensus of plausible ML ingroup tree set on SMSA
- SMSA-CE: Extended majority rule consensus of plausible ML ingroup tree set on SMSA
- SMSA-P: Plausible ML tree set for SMSA
- SMSA-B: Best-scoring ML ingroup tree inferred on SMSA

The notation for inferred trees is analogous for the FMSAO and SMSAO datasets that do also not include the outgroups in the tree search step. We will use the MSA versions of FMSAO and SMSAO that do include the outgroup sequences in a separate step for assessing outgroup rooting.

### Tree Thinning

At the time of writing, more SARS-CoV-2 sequences become available on a daily basis, while the available phylogenetic signal is already comparatively weak. To this end, it can be desirable to reduce the number of sequences we use for phylogenetic analysis in a reasonable way. We call this reduction ‘thinning’ of a phylogenetic tree. The problem of thinning or clustering sequences on taxon-rich trees is not new in virus phylogenetics (e.g., see (34,35)). To thin a given tree or input MSA prior to phylogenetic analysis one has two options. First, one can use biologically reasonable *ad hoc* criteria as we already apply them here by removing singleton and duplicate sequences, or as deployed by nextstrain that removes sequences that are below a certain length threshold. In addition, nextstrain randomly sub-samples sequences within predefined geospatial groups, to yield the inference process more computationally tractable. Second, one can deploy an inferred comprehensive phylogeny to guide the thinning process. We present and make available one MSA-based and one phylogeny-based thinning method in the following.

The first method which we term ‘maximum entropy’ selects a given number of representative sequences from the alignment that maximize sequence diversity (as measured by their entropy). The second method which we term ‘support selection’ relies on bipartition support values to identify a subset of sequences with more stable phylogenetic signal.

The ‘maximum entropy’ method aims to represent the original MSA in *n* sequences that capture as much of its diversity as possible. The method takes as input an MSA and a target number *n* of sequences to select from that MSA. First, we select a ‘seed’ sequence by finding the sequence in the MSA that is most different from a consensus sequence computed for the entire MSA; here, we measure the sequence difference as the number of non-identical sites (i.e., the Hamming distance). We use this seed to initiate the algorithm with a sequence that is as different as possible from all others. Then, the remaining *n* − 1 sequences are iteratively added to the result sequence set. In each step, we select one sequence from the yet unselected sequences of the MSA and include it in the result sequence set. We select the new sequence such that it maximizes the average per-site entropy of the current result sequence set. Hence, in each step, we greedily maximize the diversity of the current result set, as measured by its entropy; see (36) for details on the computation. The algorithm terminates once the result set contains *n* sequences (the initial seed, and *n* − 1 sequences chosen via entropy maximization), and then outputs the result set.

The ‘support selection’ method takes as input an unrooted multifurcating consensus tree *T* (here either the SMSA-C/SMSA-CE or FMSA-C/FMSA-CE trees) with a support value associated to each internal branch/bipartition of the tree. We define the *accumulated support value* (ASV) of a tree as the sum of the support values over its internal branches. Our support selection method constructs a bifurcating tree 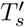 by pruning subtrees from *T* such that the ASV of 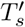 is maximized. If, for the sake of simplicity, we initially assume that *T* is rooted, we can traverse *T* in post-order: at each inner node, our algorithm selects the two children (subtrees) with the highest ASV, and calculates the ASV of the current node as the sum of the two selected children ASVs and the support value of the current bipartition. If the current node has more than two children (i.e., it is multifurcating), we prune the children that we did not select. If *T* is unrooted, we iterate over all possible inner nodes *r* of *T*, and consider each *r* as possible root of *T*. Note that each inner node *r* only constitutes a ‘virtual’ root that is required to initiate the recursion, but that the bifurcating tree 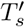 remains unrooted. In particular, *all* internal nodes of 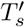 have three outgoing branches. This includes the specific virtual root *r* used for the recursion. Thus, when computing the ASV at the current virtual root *r*, we select three children subtrees instead of two. We then return the bifurcating tree 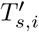 for *i* = 1 … *r* with the highest ASV.

A disadvantage of the ‘support selection’ method is that we can not control the number of taxa that we will prune. Consider, for instance, the scenario of a multifurcation with 10 children subtrees of equal size (where the size of a subtree is the number of terminal nodes in the subtree). In this case, the ‘support selection’ method will prune 8 of these subtrees.

A key question that arises is how we assess the quality of a tree thinning method and how we compare these methods against each other. In our study we consider topological stability as quality criterion. More specifically, we assess if the reduced taxon set yields a higher topological stability in terms of pairwise relative Robinson-Foulds (RF) distances (37) among the trees in the plausible tree set and a lower number of trees in the plausible tree set than a random thinning/sub-sampling of the taxa to the same number. We also assess if the thinned trees exhibit higher topological stability than the full trees on the comprehensive alignments that include *all* taxa.

### Outgroup Rooting with EPA-NG

To place the pangolin and bat outgroups onto our inferred ingroup phylogenies (more specifically, the respective plausible tree sets: FMSA-P, SMSA-P), we use our evolutionary placement algorithm EPA-NG (38) as it allows to place an arbitrary number of candidate outgroup sequences onto a given phylogeny *after* the ingroup inference. For each branch of the ingroup EPA-NG computes an outgroup placement probability via a likelihood weight ratio (LWR). The LWR indicates how probable it is that the outgroup is located somewhere along a specific branch. In other words, the methods implemented in EPA-NG allow to assess outgroup placement uncertainty and can help to answer the question if the pangolin and/or bat sequences constitute appropriate outgroups for the SARS-CoV-2 phylogeny.

As input, EPA-NG requires the ingroup tree, a MSA comprising the ingroup sequences (in this context called reference tree and reference MSA, respectively), and the outgroup sequences aligned against this reference MSA. For the alignments that included the outgroups (FMSAO, SMSAO) this is straightforward, as the outgroups are already aligned to the reference. To also assess the impact the specific MSA method has on the outgroup placements, we also deployed HMMER (39) to align the outgroups to the corresponding ingroup alignments (FMSA, SMSA) yielding two additional reference MSAs which we denote by FMSAO-HMMER and SMSAO-HMMER.

Furthermore, EPA-NG requires a strictly bifurcating input tree. This poses a challenge for trees containing multifurcations (FMSA-C, SMSA-C), as there exist alternative approaches to making such trees strictly bifurcating. To obtain a better understanding of the behavior of outgroup placements with EPA-NG in such cases, we performed placements on all trees in the respective plausible tree sets (FMSA-P, SMSA-P) that form the basis for the respective consensus trees. We evaluate the appropriateness of the bat and pangolin outgroups for each tree in the respective plausible tree sets individually by identifying the LWR of the best placement, as well as the entropy of the LWR distribution for a given outgroup.

We calculate the entropy as

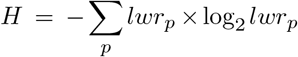

where *lwr_p_* denotes the LWR of an individual placement of an outgroup sequence on a branch *p*, for all placements calculated by EPA-NG.

We then summarize these values for the entire plausible tree set using the mean and standard deviation.

We present these statistics separately for each of the four reference MSA versions (FMSAO, SMSAO, FMSAO-HMMER, SMSAO-HMMER) in the respective results section on rooting the virus phylogeny.

### Rooting with RootDigger

To further assess the uncertainty of the root location via a mathematically distinct approach, we also performed analyses with our RootDigger tool (40). RootDigger computes the likelihood of placing a root on every branch of an existing, strictly bifurcating tree topology using a non-reversible model of nucleotide substitution. This also allows to quantify root placement uncertainty, again, by calculating LWRs for each possible root placement in terms of root placement probabilities. As RootDigger represents an alternative to outgroup rooting (albeit outgroup rooting and RootDigger rootings agree on 50% of empirical datasets tested (40)), we only executed RootDigger on the FMSA-P and SMSA-P tree sets. As the input trees are large in terms of number of taxa, we also parallelized RootDigger using MPI (Message Passing Interface) to maximize throughput.

We applied RootDigger to evaluate the root placement uncertainty for 5% of the trees with the highest likelihood scores in the respective FMSA-P and SMSA-P tree sets. We only execute RootDigger on 5% of the best trees in the plausible tree sets due to excessive runtime requirements. As for EPA-NG, we subsequently calculate analogous summary statistics for the root placement probability distributions over the respective selected plausible trees.

### Species Delimitation with mPTP

The mPTP (41) tool implements a method for molecular species delimitation on given, rooted and strictly bifurcating phylogenies of barcoding or other marker genes via so-called multi-rate Poisson Tree Processes.

As such, it exclusively relies on the tree topology and the associated branch lengths to infer a maximum likelihood based delimitation. It can also sample candidate delimitations via a MCMC procedure. In a recent study we have shown that mPTP can be successfully deployed to hepatitis type *B* and type *C* virus phylogenies for classifying sub-types (42). To this end, we applied mPTP to all trees in FMSA-P and SMSA-P.

In general, mPTP requires a *rooted* strictly bifurcating input tree. If the tree is not rooted, mPTP will by default place a root in the middle of the longest branch of the tree. If one does not trust this rooting approach, one can also execute mPTP on all distinct possible rootings of a given unrooted tree. This is not an option provided by mPTP but needs to be explicitly scripted. To assess the impact of root selection on the species delimitation, we executed mPTP maximum likelihood delimitations with all possible roots on all trees in the respective plausible tree sets (i.e., thousands of mPTP runs per plausible tree topology) via an appropriate script. We also executed delimitation runs with mPTP by rooting on the longest branch (i.e., one mPTP run per plausible tree) using the maximum likelihood and MCMC procedures.

## Difficulties

Based on our experience with the development of likelihood-based phylogenetic inference tools, we expected the phylogenetic analyses to be numerically challenging because of the large number of highly similar sequences. In fact, the dataset has a structure that is more similar to a typical population genetics dataset than a phylogenetic dataset because it exhibits, for instance, a high proportion of invariable sites. To this end, the key difficulty we expected were numerical instabilities associated with the short branch lengths.

We obtained the following results on numerical (in)stability with an earlier snapshot of the dataset.

### Impact of the minimum branch length parameter setting

We initially tested the impact of the minimum allowed branch length parameter setting on RAxML-NG and IQ-Tree log likelihood scores. For this, we used *the* best-known comprehensive tree (i.e., FMSA-B), the Γ model of rate heterogeneity, and set `blmax` (i.e., the maximum branch length) as well as `lh`_*ϵ*_ (i.e., the likelihood difference between successive numerical optimization steps used for stopping the optimization) to their default values (see Table 1).

With all these parameters fixed, we then maximized the ML score (by optimizing branch lengths and the remaining model parameters) of the fixed tree under different minimum branch length settings. We show the results of this experiment in Table 1. We observe that the minimum allowed branch length setting has a substantial impact on the resulting log-likelihood score. For instance, the default setting for IQ-Tree (`blmin` = 1e–6) yields a log-likelihood score that is 35 log likelihood units worse than for `blmin` = 1e–7. These small differences in log likelihood scores can have detrimental impact, for instance, when assessing the significance of obtained topologies via the Shimodaira-Hasegawa (43) likelihood ratio test. Initial experiments under `GTR+FO+R4` yielded an analogous, yet even more distorted ordering of log likelihood scores (data not shown).

Based on the results presented in Table 1 we therefore conducted all log likelihood score calculations with IQ-Tree for determining the plausible tree set via the statistical significance tests as well as all tree searches with RAxML-NG using a minimum branch length setting of 1e–9.

### Unreliable Scores under the Free Rates Model

Runs of ModelTest-NG (27) that we conducted on earlier snapshots of the dataset suggested that the best fit model is GTR with an ML estimate of the base frequencies and a free rates model of rate heterogeneity.

It is common knowledge among developers of ML inference programs that the numerical optimization of the free rates model is difficult and that the optimization can become stuck in local optima.

To this end, we performed model parameter and branch length re-optimization on the 100 fixed ML trees inferred with RAxML-NG on an earlier snapshot of the data using RAxML-NG and IQ-Tree. Besides the branch lengths, the model parameters were also re-optimized independently and from scratch for each ML tree (as opposed to conducting this once for the entire tree set). We conducted ML re-optimization under the GTR model with a ML estimate of the base frequencies and the four free rates that accommodate rate heterogeneity (`GTR+FO+R4`). To ensure that IQ-Tree optimizes all model parameters independently from scratch for each tree, it has to be invoked separately for each tree. In contrast to this, we can pass the entire tree set to RAxML-NG for evaluation. To assess log likelihood score discrepancies, we calculated the Spearman rank correlation on the resulting log likelihood scores obtained from RAxML-NG and IQ-Tree for the 100 ML trees. We show the log likelihood score correlation in Figure 1.

**Fig. 1.**
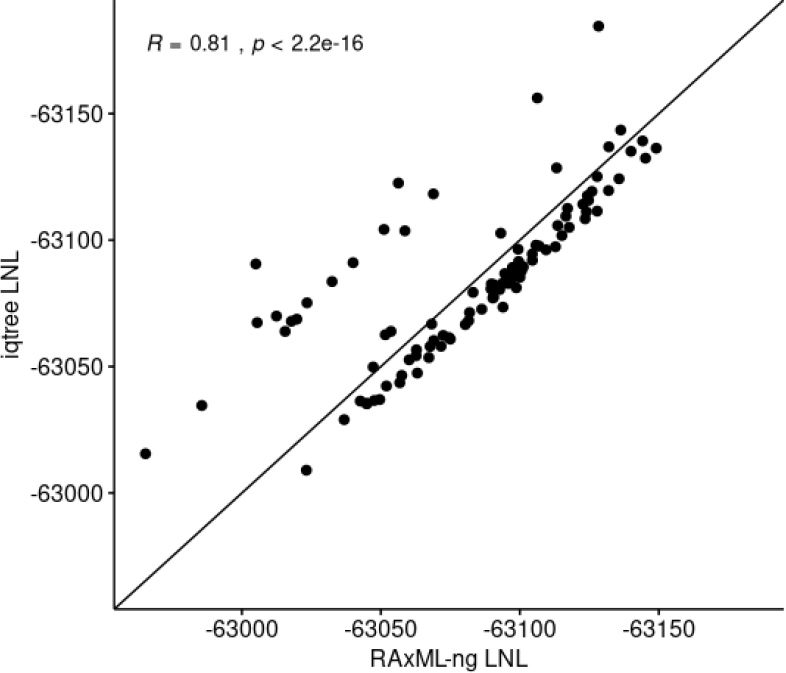
Spearman rank correlation of RAxML-NG and IQ-Tree log likelihood scores under the free rates model on a set of 100 fixed tree topologies.

The obtained correlation of merely 0.81 between the IQ-Tree and RAxML-NG scores for *exactly* identical tree topologies under *exactly* the same model of evolution indicates that the free rates model should not be used for the SARS-CoV-2 dataset. In addition, the numerical stability of the model should be reviewed in general on a broader benchmark of empirical datasets.

To ensure that this behavior is model-specific and not dataset-specific we also conducted an analogous test under the Γ model of rate heterogeneity with 4 discrete rates (`GTR+FO+G`). The corresponding log likelihood scores calculated by RAxML-NG and IQ-Tree are shown in Figure 2. With a correlation of 1.0 and based on an exactly identical likelihood-based ordering of trees scored by RAxML-NG and IQ-Tree, we conclude that the Γ model of rate heterogeneity should be used for the SARS-CoV-2 dataset.

**Fig. 2.**
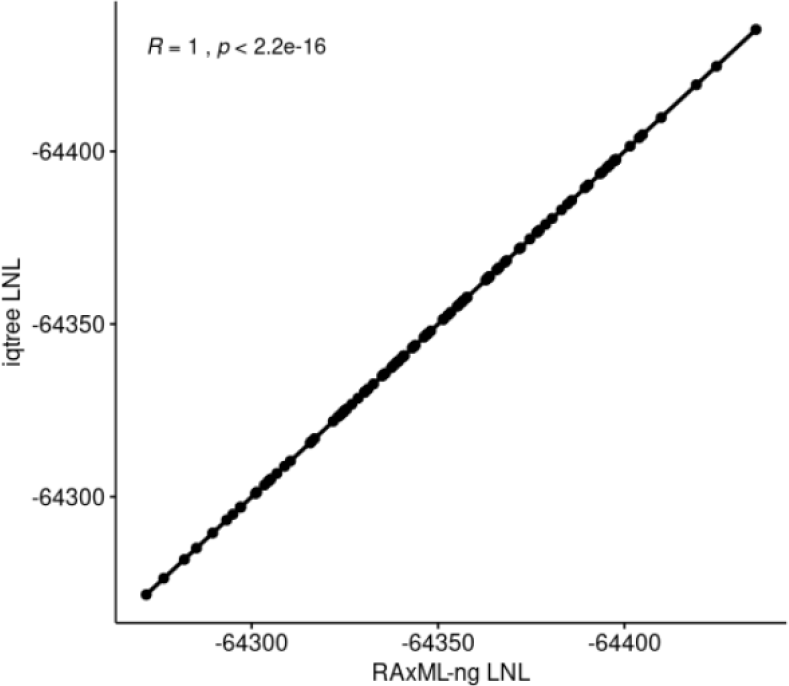
Spearman rank correlation of RAxML-NG and IQ-Tree log likelihood scores under the Γ model of rate heterogeneity on the same set of fixed tree topologies as in Figure 1

## Results

### Tree Inference

To discuss the results of the tree inferences we initially need to define the resolution ratio of the consensus trees we inferred from the plausible tree sets. Let *T* be a multifurcating tree, *B*(*T*) the number of internal bifurcating nodes, and *L*(*T*) the number of leaves. We define the *resolution ratio* of *T* as

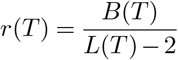

This ratio measures to which degree a tree is resolved. For instance, *r*(*T*) is equal to 0.0 for a star topology and equal to 1.0 for a fully bifurcating (fully resolved) tree.

Overall, we computed 6 metrics on the distinct trees and tree sets inferred on the four different MSA versions. We summarize these metrics in Table 2. For each metric, we obtained approximately identical values for all four MSA versions. Thus, removing singletons does not appear to improve the stability of the inferred trees, albeit the reduced taxon set can facilitate visualization and interpretation.

**Table 2.** Metrics for assessing the quality of the tree inference conducted on the four distinct MSA versions (FMSA, FMSAO, SMSA, SMSAO). *ML trees RF* is the average relative RF distance between all 100 inferred ML trees. *Search RF* is the average relative RF distance between the parsimony starting trees and the final ML trees of the respective tree searches on these starting trees. *Plausible trees* represents the number of trees (out of 100) in the plausible trees set. *MR* and *MRE resolutions* are the resolution ratios (see definition in the text) of the MR and MRE trees computed on the plausible tree sets.

| Metric | FMSA | FMSAO | SMSA | SMSAO |
| --- | --- | --- | --- | --- |
| Taxa | 4,869 | 4,871 | 2,888 | 2,904 |
| ML trees RF | 0.783 | 0.783 | 0.775 | 0.783 |
| Search RF | 0.112 | 0.112 | 0.128 | 0.119 |
| Plausible trees | 76 | 75 | 74 | 76 |
| MR res. $r(T)$ | 0.129 | 0.131 | 0.155 | 0.147 |
| MRE res. $r(T)$ | 0.706 | 0.701 | 0.699 | 0.680 |

Moreover, for all MSA versions, we inferred 100 *distinct* topologies from the 100 ML searches (i.e., the signal was so weak that we did not recover a single tree topology twice). Furthermore, the ML tree topologies per MSA are highly different among each other with an average pair-wise relative RF distance of approximately 78%.

In contrast, the relative RF distance between the respective parsimony starting trees and the corresponding final ML trees of individual searches is comparatively low (ranging between 0.11 to 0.13). This indicates that every ML tree search quickly reaches a local maximum and that all MSA versions induce a high number of local maxima. In general, approximately 75 out of 100 inferred ML trees per MSA end up in the respective plausible tree sets. This shows that it is difficult, if not impossible, to distinguish among the topologically highly diverse ML trees from 100 searches via statistical significance tests. Hence, it does not appear reasonable to represent the results in the form of a single ML tree, as 75% of the inferred trees are indistinguishable.

We further found that the majority rule consensus trees (FMSA-C and SMSA-C) we computed from the plausible tree sets are poorly resolved and only contain but a few bifurcating internal nodes. The extended majority rule trees (FMSA-CE and SMSA-CE), which attempt to construct bifurcating tree topologies via a greedy heuristic strategy (note that constructing the optimal MRE consensus with maximum support is NP-hard) still contain a high number of multifurcating nodes. Nonetheless, they do show an improved degree of resolution (*r*(*T*)) by a factor of 4 to 5 compared to the MR trees which simplifies their visual interpretation.

Overall, we find that the ML tree topologies in the plausible tree sets are topologically divergent which substantiates our claims that the dataset is hard to analyze as it exhibits a weak phylogenetic signal and a rugged likelihood surface. Our experiments also show that results of phylogenetic analyses of these data can and should not be represented via a single ML tree. Our findings contradict a recent study (44) that finds that there is sufficient phylogenetic signal in the data. This study relies on the so-called likelihood mapping technique that only evaluates quartets (subsets of four sequences) to quantify the signal. Therefore, the aforementioned numerical issues associated with a full tree search on a comprehensive MSA did not become apparent. Finally, we observe that the specific MSA version used does not have any notable effect on the resulting tree set.

In the following, we briefly discuss the virological conclusions that we can draw by example of the FMSAO-CE tree.

The FMSAO-CE consensus tree (see Figure 3) suggests that the clades that occur frequently in the respective plausible tree set (> 75%) consist predominantly of SARS-CoV-2 sequences sampled from the same geographic area or neighbouring countries. Specifically, we find large monophyletic clusters from a single country or geographic area such as the USA, India, Hong Kong, Shanghai, Korea, Iceland, Wales, Scotland, England, Australia, Belgium, Luxembourg, the Netherlands, France, etc. We detected the largest clusters for the USA and England. Moreover, we observe clusters including sequences from neighbouring geographic areas, for example, Wales – England – Scotland, Luxembourg – Belgium, Belgium – Netherlands, Scotland – Iceland. We observe two additional characteristic patterns: (i) clusters where the majority of sequences are from a single country and (ii) clades including viral sequences sampled from diverse locations. The type (i) clusters include Sweden – Wales – England, Australia – USA, England – Australia, England – Russia – Australia – Hungary – the Netherlands – USA. The more diverse type (ii) clusters are smaller in size and comprise viral sequences sampled at diverse locations.

**Fig. 3.**
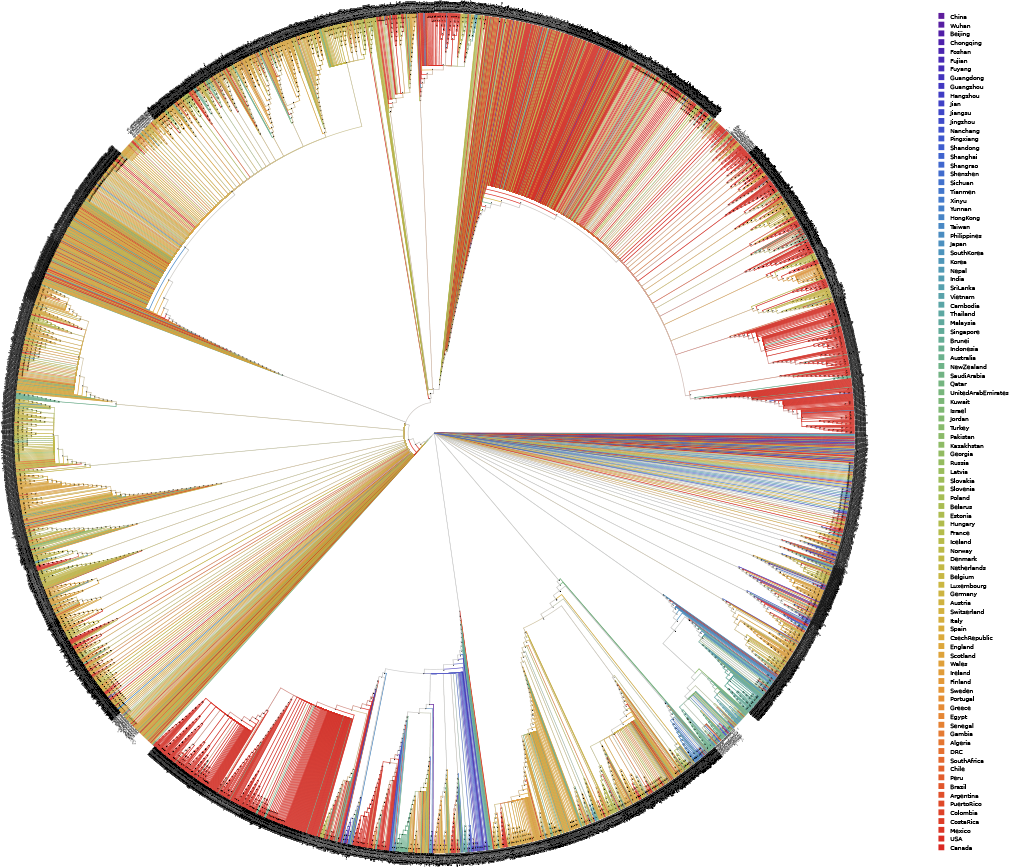
Extended majority rule consensus tree (FMSAO-CE) of the plausible tree set of the FMSAO alignment. We coloured the tree by the country of origin of each sequence.

The observed patterns suggest that clustering and thus spread occurs mainly according to geographic location. This finding is compatible with the diseases spread through respiratory particles, but also across different countries and remote locations following the patterns of human mobility. The results of nextstrain analyses, where major clades were detected for the USA and other regions, but also a considerable number of cross border transmissions was reported, support our findings. Notably, our and other analyses are limited by the available data sampling. The lack of large monophyletic clusters for several geographic areas is probably due to the limited availability of data from the respective countries.

### Tree Thinning

For the sake of simplicity, we executed both thinning methods only on FMSA and SMSA. We executed the support selection thinning method on the respective MRE consensus trees (FMSA-CE, SMSA-CE) instead of the MR consensus tree as it yielded a thinned tree comprising an order of magnitude less taxa. As the MR consensus comprised too many multifurcations it yielded too small thinned alignments (comprising less than 50 taxa) that were not apt for biological interpretation. As maximum entropy thinning does not require a tree as an input (see description of the method) we executed it directly on the FMSA and SMSA alignments. Finally, we also executed a naïve thinning by randomly removing a given number of sequences from the initial SMSA and FMSA alignments, to assess if our two thinning approaches perform better than random thinning.

For improved clarity, we introduced the following notations for the different thinned alignments we computed:

- F-SST: alignment obtained from the FMSA alignment using support selection thinning.
- F-MET: alignment obtained from the FMSA alignment using maximum entropy thinning.
- F-RAND: alignment obtained from the FMSA alignment using random thinning.
- FMSA-SS-P: plausible tree set of F-SST.
- S-SST: alignment obtained from the SMSA alignment using support selection thinning.
- S-MET: alignment obtained from the SMSA alignment using maximum entropy thinning.
- S-RAND: alignment obtained from the SMSA alignment using random thinning.
- SMSA-SS-P: plausible tree set of S-SST.

We calculated the same quality metrics as used for the tree inferences on the comprehensive non-thinned MSAs for the thinned MSAs in Table 3. We find that the stability of tree inferences is slightly improved by support selection thinning and maximum entropy thinning. The average relative RF distance between all 100 inferred ML trees decreases on all four alignment versions from approximately 0.78 down to 0.67 using support selection thinning and down to 0.63 via maximum entropy thinning. The resolution (*r*(*T*)) of the consensi improves for support selection thinning as well as maximum entropy thinning. The better MRE resolution of support selection thinning versus maximum entropy thinning is due to the design of the support selection algorithm that uses the MRE on the comprehensive tree as an input. In other words, the method works as intended. Further, we can reduce the size of the plausible tree set by approximately 40 – 50% with both approaches. Nonetheless, the substantial reduction in the number of taxa by these thinning approaches does not alleviate the problem of weak signal and multiple ML optima. Finally, we find that support selection thinning and maximum entropy thinning perform consistently better than random thinning.

**Table 3.** Metrics for the *thinned* alignment versions. *Taxa* is the number of taxa in the alignment. *ML trees RF* is the average relative RF distance between all 100 inferred ML trees. *Search RF* is the average relative RF distance between the parsimony starting trees and the final ML trees of the respective tree searches on these starting trees. *Plausible trees* represents the number of trees (out of 100) in the plausible tree sets. *MR* and *MRE resolutions* are the resolution ratios (see definition in the text) of the MR and MRE trees computed on the plausible tree sets.

| Metric | F-SST | F-MET | F-RAND | S-SST | S-MET | S-RAND |
| --- | --- | --- | --- | --- | --- | --- |
| Taxa | 912 | 912 | 912 | 434 | 434 | 434 |
| ML trees RF | 0.67 | 0.66 | 0.77 | 0.68 | 0.63 | 0.79 |
| Search RF | 0.19 | 0.20 | 0.15 | 0.21 | 0.21 | 0.18 |
| Plausible trees | 39 | 45 | 73 | 31 | 47 | 59 |
| MR resolution | 0.166 | 0.218 | 0.144 | 0.164 | 0.245 | 0.141 |
| MRE resolution | 0.918 | 0.842 | 0.72 | 0.912 | 0.85 | 0.72 |

### Rooting

In Tables 4 and 5 we present the results for the outgroup rooting analyses using EPA-NG for the pangolin and bat outgroup sequences, respectively.

**Table 4.** EPA-NG root placement probability and entropy statistics for the pangolin outgroup sequence over all trees in the respective plausible tree sets for distinct MSA versions. Highlighted in bold is the highest confidence signal, which is the only among all tested datasets to reach above 0.04 mean LWR.

| Alignment | Max LWR |  | LWR Entropy |  |
| --- | --- | --- | --- | --- |
|  | Mean | SD | Mean | SD |
| FMSAO | 0.033 | 0.001 | 5.332 | 0.010 |
| FMSAO-HMMER | 0.034 | 0.000 | 5.325 | 0.010 |
| SMSAO | **0.647** | 0.010 | **2.074** | 0.046 |
| SMSAO-HMMER | 0.001 | 0.000 | 5.634 | 0.000 |

**Table 5.** EPA-NG root placement probability and entropy statistics for the bat outgroup sequence over all trees in the respective plausible tree sets for distinct MSA versions.

| Alignment | Max LWR |  | LWR Entropy |  |
| --- | --- | --- | --- | --- |
|  | Mean | SD | Mean | SD |
| FMSAO | 0.037 | 0.001 | 5.437 | 0.009 |
| FMSAO-HMMER | 0.037 | 0.001 | 5.438 | 0.008 |
| SMSAO | 0.025 | 0.001 | 5.378 | 0.006 |
| SMSAO-HMMER | 0.004 | 0.000 | 5.546 | 0.013 |

We present the mean placement probability (likelihood weight) and its standard deviation for the most likely placements of all trees contained in the respective plausible tree sets obtained for the comprehensive MSAs. Remember that FMSAO and SMSAO stand for alignments conducted with MAFFT *including* the outgroups, whereas FMSAO-HMMER and SMSAO-HMMER represent the ingroup MSAs (*excluding* the outgroups) to which we subsequently aligned the outgroup sequences via `hmmalign` in a separate step. We did this to assess the potential impact of the alignment procedure onto the placement result. Note that, a mean placement probability value of 0.033 represents a placement probability of the outgroup onto the most likely branch of the reference phylogeny amounting to 3.3%. To further characterize the LWR distribution over the branches of the tree, we computed the mean and standard deviation of the entropy, calculated across each LWR distribution of the outgroup on a given tree (see Subsection: ‘Outgroup Rooting with EPA-NG’).

As Tables 4 and 5 show, support for an outgroup rooting using either the bat or the pangolin sequence was generally low (< 0.04). The only exception is a possible well-supported pangolin-based rooting of the plausible trees in the SMSAO dataset. This is surprising, as, with the exception of a small fragment in the spike protein, the pangolin is more divergent from the ingroup than the bat. For this specific alignment, EPA-NG yielded a well-supported placement of the pangolin outgroup for all plausible trees, yet always residing on the terminal branch leading to sequence `EPI_ISL_411956` (GI-SAID accession). While this sequence is among the early sequences of the pandemic, a placement of the root on the branch leading to it, does not appear to be epidemiologically plausible. After pruning `EPI_ISL_411956` from SMSAO-B and SMSAO and subsequently re-calculating the pangolin placement, the placement confidence was lower (0.176). In addition, the new placement location with 0.176 support is located at a large distance to the initial highly supported location of `EPI_ISL_411956`. The path length, in terms of inner nodes along the tree, between the initial and the new placement location amounts to 105 inner nodes.

As we suspected that the confident placement of the pangolin constitutes an artifact of the alignment process, we repeated the MSA procedure under distinct settings. Initially, we removed the bat outgroup from the initial set of unaligned sequences. This yielded an increased LWR (Mean 0.87, STD 0.002) for placing pangolin on the branch leading to `EPI_ISL_411956`). Second, we added two additional bat outgroup sequences (`MG772933` and `MG772934`) to the unaligned sequence set. While this lowered the LWR of the pangolin placement (Mean 0.206, STD 0.01, again located on the branch leading to `EPI_ISL_411956`), the signal was still considerably stronger than for all other outgroups and MSA versions. A visual inspection of the SMSAO alignment to identify the reasons for the strong pangolin placement signal was inconclusive. The same holds true for an inspection of the per-site log likelihood values for pangolin placements into distinct branches (including the highly supported one) of the corresponding best ML tree (SMSAO-B).

Overall, despite our efforts, we were not able to disentangle the reasons behind this strong, yet epidemiologically implausible placement of the pangolin. The additional experiments we conducted using alternative outgroups indicate that this is potentially due to an alignment artifact in just one out of the four alignment versions we scrutinized. Hence, not only the tree inference itself, but also the alignment strategy used can impact the results of phylogenetic post-analyses of SARS-CoV-2.

We present our results for the root placement certainty as calculated with the RootDigger tool on the ingroup phylogeny in Table 6. We executed RootDigger searches on the 4 plausible tree sets inferred on the FMSA, SMSA, FMSA-SS, SMSASS (thinned FMSA and SMSA alignments with support selection thinning, see previous section). As mentioned above, we performed RootDigger analyses only on the top 5% of the trees in the respective plausible tree sets due to excessive runtimes. This resulted in performing rooting analysis on 8 trees from FMSA-P, 4 trees from FMSAN-SS-P, 8 trees from SMSA-P, and 4 trees for SMSA-SS-P. We used the same method to calculate the LWR entropy of RootDigger root placements as for the EPA-NG results above.

**Table 6.** Results of RootDigger analysis for different MSA versions. Because of excessive runtimes, for every dataset, we only analyzed the 5% of trees with the highest likelihood with RootDigger in exhaustive mode. To further summarize the results, we also compute the entropy of the LWR distributions for each resulting tree and report the average for each dataset. The results are averages over the included plausible trees.

| Alignment | Max LWR |  | LWR Entropy |  |
| --- | --- | --- | --- | --- |
|  | Mean | SD | Mean | SD |
| FMSA | 0.240 | 0.006 | 8.053 | 0.059 |
| FMSA-SS | 0.041 | 0.001 | 7.872 | 0.036 |
| SMSA | **0.613** | 0.038 | **4.279** | 0.474 |
| SMSA-SS | 0.101 | 0.002 | 7.327 | 0.023 |

While the root inferences on the thinned MSAs do not yield strong signal for any particular root placement, this is not the case for the original alignments. In particular, we obtain a strong average (over the 8 trees with the highest likelihood score) root placement signal for a specific root on the SMSA alignment which we discuss in further detail below.

To this end, we visually inspected the 8 rooted trees for SMSA as inferred with RootDigger. For 2 out of 8 trees (tree 1 and 2, trees are labelled 0 – 7) we observed an epidemiologically plausible root placement, since among the sequences which cluster close to the inferred root, there are several from Wuhan and other Asian areas sampled during the early phase of the pandemic. We show the respective rooted maximum likelihood tree number 2 colored by geographic regions in Figure 4. Nonetheless, we obtained such a virologically plausible root placement with RootDigger only for 25% of the rooted trees and only for one out of 4 alignment versions. Hence, the plausibility of the root placement heavily depends on the selected MSA version as well as selected tree from the plausible tree set and the results can not be generalized. The same holds for the outgroup placements with EPA-NG. Here, while we do again observe a relatively strong signal only on one out of 4 aligment versions, the outgroup placement location does not appear to be virologically plausible. Thus, we conclude that the root of the SARS-CoV-2 phylogeny can not be reliably determined via the methods we have applied here. An independent study on root placement using distinct computational methods comes to analogous conclusions (22).

**Fig. 4.**
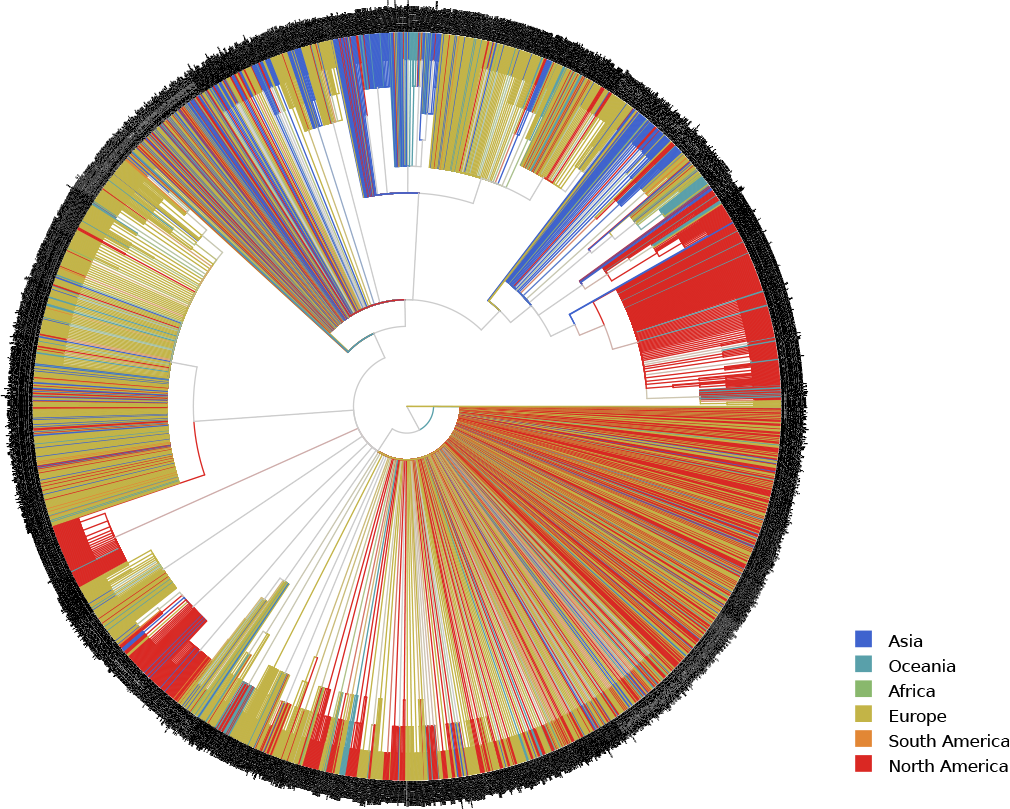
Rooted SMSA Maximum Likelihood tree number 2. We colour the tree by geographic regions and root it via RootDigger using a non-reversible model of nucleotide substitution. The tree inference randomly resolved multifurcations by introducing branches of length zero. For visualization purposes, we collapsed these branches, hence yielding a multifurcating tree again.

### mPTP

The mPTP runs on all plausible tree sets using the longest branch rooting option and either the ML delimitation or the MCMC delimitation procedures yielded a species count of 1. This means that mPTP in default mode can not distinguish if there is 1 species or if there are *n* species, where *n* is the number of taxa in the given phylogeny.

The mPTP runs that explored all possible rootings on all plausible tree sets under the ML delimitation option exhibit a large variance in results.

We show a representative histogram for the SMSA-P set of plausible trees in Figure 5 for the median number of delimited species over all possible rootings per plausible tree. The results on the remaining datasets were analogous (data not shown). The maximum number of delimited species for all rootings per tree in SMSA-P ranged between 198 and 781 with a flat distribution (i.e., two identical maximum species counts appeared 9 times, and three identical ones only once). The minimum number of delimited species was 1 for all trees in SMSA-P.

**Fig. 5.**
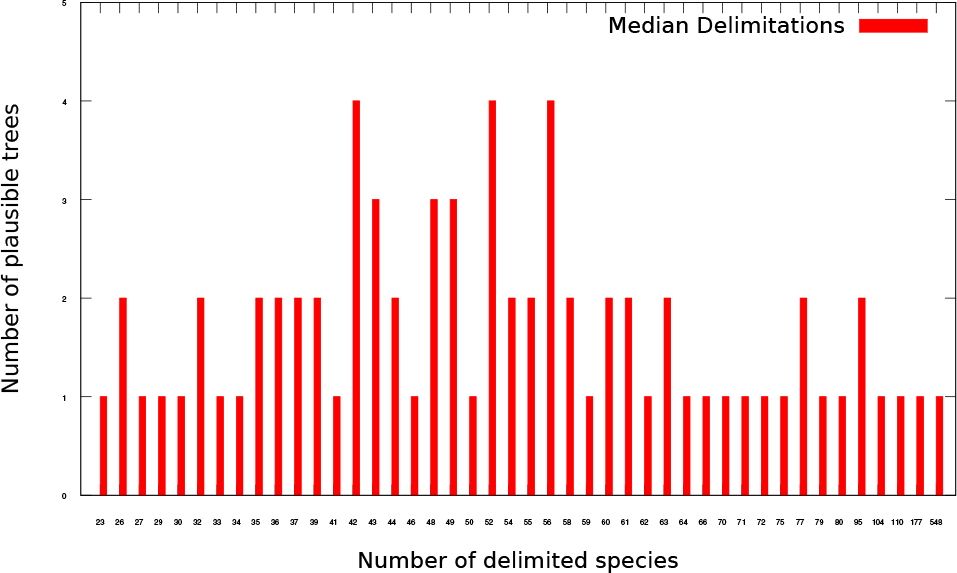
Median number of delimited species over all possible rootings per plausible tree in SMSA-P.

We hence conclude that mPTP can not be used to delimit distinct sub-classes of the virus as the default mode (rooting at the longest branch) consistently yielded inconclusive delimitations. Further, as shown by our experiments that evaluate all possible rootings, the number of delimited species exhibits a large variance as a function of the root position and is hence also inconclusive. Given that, the trees can, in general, not be reliably rooted, we conclude that we can not delimit/classify the viral sequences using mPTP. Other authors have also put into question our ability to identify distinct virus types using alternative computational methods (45).

## Discussion

We studied the intrinsic difficulties of inferring and post-processing phylogenetic trees on the May 5 snapshot of the available whole genome data for SARS-CoV-2. To quantify the impact of distinct filtering and alignment strategies, we use four different alignment versions throughout our analyses.

We find that the tree search task *per se* is difficult due to the rugged likelihood surface that exhibits a multitude of local optima. We can not distinguish among the majority of these local optima via standard statistical significance tests and observe large pair-wise topological differences that exceed 70%. We therefore suggest that instead of using and displaying a single tree one should compute summary statistics on a ’plausible tree set’ that comprises the indistinguishable local maxima of the tree search space that were found by the respective search algorithm.

While using a ML approach to infer trees, our post-analysis strategy rather follows a Bayesian paradigm. Thus, the question arises if one could use Bayesian inference via MCMC methods directly. Because of the size of the datasets, their lack of signal, and the large number of taxa we expect MCMC analyses to require excessive runtimes to reach and explore some of the peaks we identified with the more targeted ML searches. In addition, our experience, based on user interactions, is that MCMC analyses are more difficult to properly set up and interpret than ML analyses, especially on such a challenging dataset that requires a profound understanding of the underlying methods.

To this end, the current common practice of inferring SARS-CoV-2 phylogenies under the default search parameters of standard ML inference tools corresponds to randomly picking (without potentially being aware of it) a tree from the plausible tree set. Due to the large topological variations, the respective conclusions that we draw can constitute a product of pure chance.

Beyond this, we also identified substantial numerical issues pertaining to the optimization of branch lengths and the rates in the free rates model of rate heterogeneity. Branch length optimization is problematic because the sequences are highly similar and, as a consequence, the branch lengths are short. Thus, assessing the effect of the minimum branch length setting in ML inference tools on the results constitutes a necessary prerequisite for conducting thorough phylogenetic analyses of these data. The issues associated with the parameter optimization in the free rates model are likely to not only occur on difficult datasets, but we believe that they become more prevalent on such.

We also address the problem of reducing the size of the plausible tree sets in the hope that a reduction in size will induce a decrease of average pair-wise RF distances and an increase of consensus tree resolution, thereby simplifying the interpretation and post-analyses. In addition, we can also interpret a reduction of the plausible tree set size as an indicator of stronger signal. To this end, we introduce and test two novel tree thinning algorithms that strive to maximize the entropy and support of the thinned alignments and respective trees. While these algorithms do reduce the size of the plausible tree sets and perform better than random thinning, the plausible tree sets still remain comparatively large (comprising approximately 40 out of 100 ML trees) and diverse (average pair-wise topological RF distance among the plausible trees slightly below 70%).

Overall, we believe that using an extended majority rule consensus tree inferred on the plausible tree sets represents a reasonable approach to carefully interpreting the results by taking into account the ruggedness of the tree search space. For certain epidemiological assessments, it will suffice if the branching order near the tips of the phylogeny is well resolved.

With respect to post-analyses, we find that rooting the trees either via outgroup placement or by using non-reversible models of evolution does *not* yield a clear root position. Obtaining an epidemiologically reasonable root with some statistical support appears to be a matter of chance: it depends on the specific tree topology used for conducting the rooting analysis, which we selected from the plausible tree set inferred from one specific alignment. With respect to outgroup placement, the single strong, yet epidemiologically implausible signal was also observed on one specific alignment version only. We can not draw general, nor confident conclusions about the position of the root using the two mathematically highly distinct approaches that we have deployed here. This confirms analogous independent findings (22).

Finally, we find that distinct viral sub-classes can not be identified by executing our mPTP tool for molecular species delimitation on all trees in the respective plausible tree sets of all four alignment versions.

## Conclusions

Phylogenetic analysis of SARS-CoV-2 data is challenging due to numerical difficulties and the rugged likelihood surface. Therefore, we strongly advocate against naïvely using the default parameters of common ML programs to just infer ‘a tree’ and using this single tree for epidemiological interpretation or any type of post-analysis.

We are also skeptical about the utility of computing bootstrap support values, as the datasets as such only contain a low number of distinct site patterns while containing thousands of taxa. One should preferably invest computational effort to more exhaustively sample the rugged likelihood surface which already constitutes a source of large topological variability.

As the phylogenetic signal is weak, we suggest using a plausible tree set comprising all ML trees from independent tree searches that we can not distinguish from each other via the standard phylogenetic significance tests, but that does adequately represent the rugged tree search space.

In analogy to summarizing results from Bayesian tree inferences, we suggest to use this plausible tree set for computing summary statistics on the trees such as the MR or MRE consensi. Also, we should conduct and summarize all potential post-analyses on such plausible tree sets to better capture topological uncertainty and circumvent potential misinterpretations that can be caused by randomly picking a tree from the plausible tree set.

To this end, we believe that we need to develop novel methods that can automatically summarize such plausible tree sets. In addition, there is a need for theoretical work on criteria to identify datasets that exhibit rugged likelihood surfaces, as the term is admittedly colloquial and vague at present. Ideally, phylogenetic inference programs should be able to identify such difficult datasets and either warn the users about it or by default conduct multiple ML searches and automatically return a plausible tree set.

## Supporting information

Gisaid acknowledgement file

## ACKNOWLEDGEMENTS

Part of this work was funded by the Klaus Tschira Foundation and through the EU IGNITE ITN project. We wish to sincerely thank all scientists and submitting laboratories (Supplementary Acknowledgement Table) involved in collecting, processing, and depositing SARS-CoV-2 sequence data as well as meta-data on GISAID.

## Supplement

**Fig. 6.**
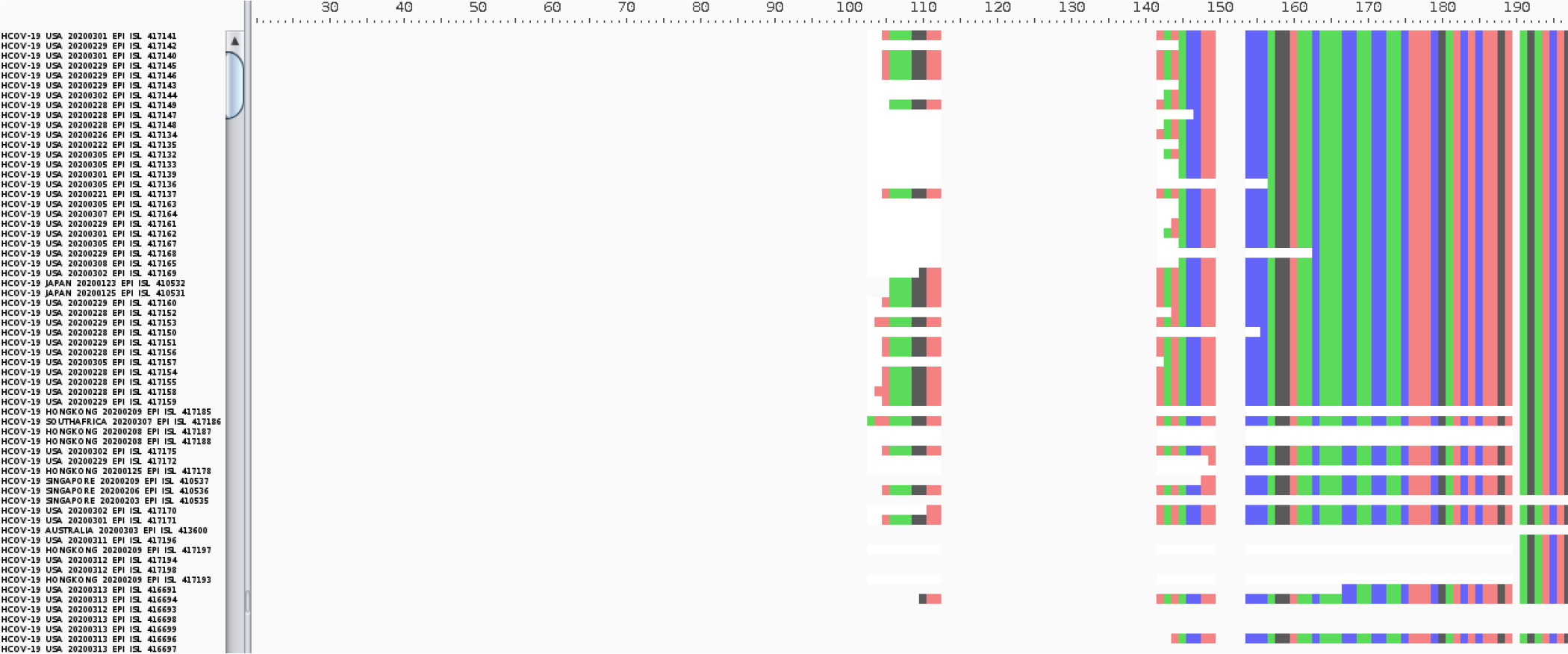
A screenshot from the Aliview alignment viewer showing the beginning of the alignment.

